# Capturing heterotachy through multi-gamma site models

**DOI:** 10.1101/018101

**Authors:** Remco Bouckaert, Peter Lockhart

## Abstract

Most methods for performing a phylogenetic analysis based on sequence alignments of gene data assume that the mechanism of evolution is constant through time. It is recognised that some sites do evolve somewhat faster than others, and this can be captured using a (gamma) rate heterogeneity model. Further, some species have shorter replication times than others, and this results in faster rates of substitution in some lineages. This feature of lineage specific rate variation can be captured to some extent, by using relaxed clock models. However, it is also clear that there are additional poorly characterised features of sequence data that can sometimes lead to extreme differences in lineage specific rates. This variation is poorly captured by constant time reversible substitution models. The significance of extreme lineage specific rate differences is that they lead both to errors in reconstructing evolutionary relationships as well as biased estimates for the age of ancestral nodes. We propose a new model that allows gamma rate heterogeneity to change on branches, thus offering a more realistic model of sequence evolution. It adds negligible computational cost to likelihood calculations. We illustrate its effectiveness with an example of green algae and land-plants. For many real world data sets, we find a much better fit with multi-gamma sites models as well as substantial differences in ancestral node date estimates.

## 1 Introduction

Systematic error is one of the major concerns of phylogenetic reconstruction, as it can mislead tree building. Heterotachy [22, 24] – the inference of lineage specific rate variation – is one problem and potential cause of systematic error. Heterotachy is present in many data sets [22]. Different underlying processes of evolution can explain heterotachy [20], one of these being that changes in intermolecular interactions over time can affect which and how many sites are free to vary [16, 15]. To some extent, the processes of nucleotide and amino acid substitution can be accommodated by covarion and gamma distributed substitution models [12, 20, 23, 28, 29]. To maintain time reversibility, these models assume that the number of sites assigned to a particular evolutionary rate class sequence remains constant through time [30]. However, this might not be a realistic biological assumption. Numerous observations suggest that the proportion of variable sites in homologues changes over larger evolutionary distances [15, 19, 21] (but see also [33] regarding sufficiency). A consequence of this form of non-stationarity, when modelling sequence evolution using a discrete gamma distribution, is that the alpha shape parameter, which describes the proportion of sites assigned to a particular rate class, will change across the underlying tree [30].

For the gamma rate heterogeneity model to be consistent, combining two samples should result in similar shape parameter estimates. However, it is well known that different sampling strategies result can in different estimates of the shape parameter [28]. This suggests that the rate heterogeneity is not constant, but changes throughout the tree, which justifies investigating models that allow such shape variation.

Here we investigate heterotachy and its impact on topology and divergence time estimates in the green plant evolutionary lineage. This is one of the oldest and most diverse branches of the tree of life. Researchers reconstructing evolutionary relationships with green plants face difficulties in phylogenetic reconstruction similar to those being faced elsewhere by others [24]. The challenge is worthwhile, as overcoming the problems will lead to insight into unresolved questions such as: What was the nature of the earliest angiosperms and land plants? How many times was land colonised from the water by “green algae”? Where did the key adaptive features for life on land come from? How many times has multicellularity arisen in the green plants? Did multicellularity ever reverse? How many times did alternation of generations and diploid-dominant life-cycles arise? While justifiable emphasis has been placed on the importance of taxon sampling, increasing attention is now being focussed on the problem of data-model fit [6]. Here, we examine the potential of multigamma and relaxed-gamma site models to improve data-model fit and in so doing improve phylogenetic inference. We describe our findings and implement these new models in the open source software BEAST and thus introduced a practical and user friendly way to model shape parameter variation throughout the tree.

## 2 Methods

First, we revisit the classic gamma rate heterogeneity model, before describing the multi-gamma model and the relaxed-gamma model.

### 2.1 Gamma site model (with invariant sites)

The likelihood of the data in an alignment *D* for tips of a tree *g* is:

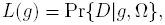

where Ω = {*Q*, *µ*} includes parameters *Q* of the substitution model and the overall rate *µ*. Consider edges 〈*i, j*〉 ϵ *R* of tree *g =* {*R*, t}, and node heights *t*_*i*_ and *t*_*j*_, and let the entry *s*_*i,k*_ be a nucleotide base at site *k* of the sequence at node *i.* As usual, we assume the sites in the alignment are independent, and the alignment represents data at the tips, but not for internal nodes. Then we sum over all possible ancestral sequences *D*_*Y*_ for assignments of internal node to get Felsenstein’s tree likelihood [11] which can be written as

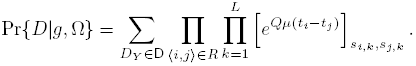

This likelihood assumes constant rates over all sites, which is not realistic in many situations. Yang et al. [34] introduced a model that assumes site rates are distributed according to a gamma distribution 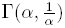 with mean one, leaving one shape parameter *α.* The likelihood can be written as

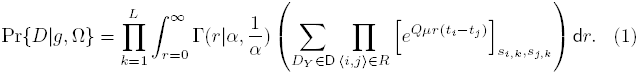

which can be efficiently calculated using Felstenstein’s peeling algorithm [11]. Computationally, it is convenient to approximate the integral by a sum over *K* categories with rates *r*_*c*_(*α*) for category *c* representing the mean of the *k*th quantile of the gamma distribution with mean 1 and shape parameter *α.* The likelihood can be calculated as a mixture of likelihoods with rates *r*_*c*_(*α*) as

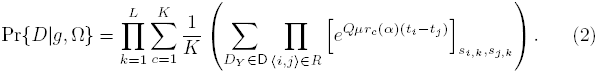

Often, there are many constant sites, and a rate of zero would fit best for these sites. Adding another category *r*_1_ = 0 with invariant sites [14, 32] for a proportion *p*_1_ of the mixture, and using the remainder for the gamma rate model with *K* categories, the likelihood becomes

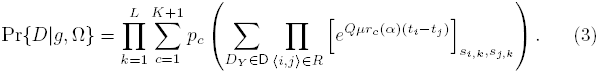

where 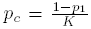 for *c >* 1 and *r*_*c*_(*α*) the mean of the *c* – 1th quantile of the gamma distribution for *c >* 1.

### 2.2 Multi-gamma site model

In the gamma rate heterogeneity model, the shape parameter *α* is assumed to be constant throughout the tree. In the *multi-gamma model* we assume that for each branch and in node *i* in the tree, we have an individual shape parameter *α*_*i*_ governing the heterogeneity of the rates. Practically, this means replacing the rate *r*_*c*_(*α*) in Equation (1) with rate *r*_*c*_(*α*_*i*_).

To distinguish this model from Yang’s model, we call the latter the *single-gamma mode* in the remainder of this paper. An obvious disadvantage is that in a tree with *n* branches, we need to estimate *n* parameters for multi-gamma model while only a single parameter needs to be estimated for the single-gamma model.

Our experience it that this is achievable in practice if there is sufficient heterotachy in the data, and it looks like a good option for maximum likelihood implementations. However, care must be taken with binary rooted trees, since the shape parameters for branches to the root appears to be unidentifiable in practice.

### 2.3 Relaxed-gamma site model

Instead of estimating the individual shape parameters, we can assume they are distributed according to a density *p*(*α*|*θ*) shared by all shape parameters and integrate them out. Pr{*D*|*g*, *Ω*} becomes

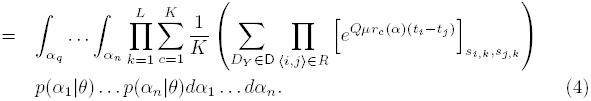

We call this model the *relaxed-gamma model* to distinguish it from the single and multi-gamma models. Note that the number of parameters in this model is just the cardinality of *θ.* If *p*(*α*|*θ*) is a log-normal or gamma distribution, there are just two parameters, which is just a single parameter more than the single-gamma model.

Unfortunately, calculating Equation (4) directly is not feasible, so we use MCMC in a Bayesian setting to approximate these integrals in a similar fashion as done for the uncorrelated relaxed model [8]. The density *p*(*α*) can be discretised into a number of quantiles. Earlier findings [18, 25] suggests that setting the number of quantiles equal to the number of branches is appropriate that number is larger than 100. For each quantile *q*, we calculate the mean value *m*_*q*_ and for each branch *i*, and we maintain an index *α*_*i*_ of a quantile for that branch. We use the following proposals

- a proposal, that randomly selects a branch and randomly selects a new value for the index *α*_*i*_ inside a window of its old value,
- a proposal that randomly selects a branch and randomly selects a new value for the index *α*_*i*_ in its range,
- a proposal that randomly selects two branches and swaps their indices,
- a proposal to scale the parameters of the density *p*(*α*|*θ*) for each of the parameters in *θ.*

### 2.4 Miscellaneous

Similar to the single gamma model with a category for invariant sites, the multigamma and relaxed gamma models can be extended to include invariant sites by generalising Equation (3) in a similar way we generalised Equation (2) by adding a category with rate zero for all branches.

*Remark:* We did not make any assumptions about the tree, so the theory applies to unrooted, rooted and rooted time trees [8], binary trees as well as polytopies and sampled ancestor trees [13].

Note that the computation complexity of the tree-likelihood – Equation (1) – adapted for the multi-gamma or relaxed gamma model using the Felstenstein’s peeling algorithm does not change, so no extra computational cost is incurred in calculating the likelihood. Of course, since the parameter space is larger for these models, longer computation is required to estimate these extra parameters.

## 3 Results

In this section, we show the effectiveness of the model using a simulation study and using randomly selected alignments from TreeBase [26], alignments from the PartitionFinder dataset repository https://github.com/roblanf/PartitionedAlignments and a case study concerning the evolutionary relationships of green plants and algae.

### 3.1 Simulation study

First, we establish how reliably the shape parameter of the single-gamma model can be estimated. The shape parameter *α* was sampled from the exponential distribution with mean 1, which is commonly used as prior for the shape parameter. Sequences of 10,000 sites were simulated under the HKY model on a tree with 3 taxa ((*A, B*), *C*) and a single-gamma model with four categories. The simulation study was performed using BEASTShell [2]. Then, an analysis under the same model was done in BEAST where *α* was estimated. Figure 1 shows the sampled *α* on the x-axis and estimated *α* on the y-axis. It shows that *α* can be estimated correctly over a large range, though it underestimates the alpha value for very large values and overestimates the alpha value for very small values. This is to be expected, since these extreme values, though rare, do not leave a large trace in the data, so the prior pulls the estimate towards the mean of 1.0. Note that smaller *α* values are estimated with lower error than higher *α* values.

**Figure 1.**
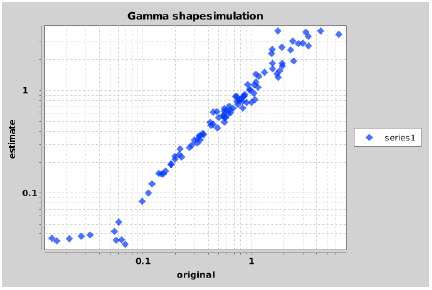
Simulated (horizontal axis) and estimated (vertical axis) values of gamma shapes when simulating with single gamma. Both axis are log-scale.

We repeated the experiment, but instead of keeping *α* constant throughout the tree, only the *α* for a single branch in taxon *A* was varied. The left of Figure 2 shows what happens when we estimated *α* using the single-gamma model; it appears that the estimated *α* is the mean of the simulated *α* values, when branch lengths are taken into account. The *α* value that was varied does not dominate the estimate. The right of Figure 2 shows *α* values for individual branches. There is some increase in the uncertainty of the estimates, but simulated values remain in the 95%HPD of estimated values, again with the exception of extreme *α* values. In conclusion, our observations show that under the simulated conditions the multi-gamma model can recover shape parameters.

**Figure 2.**
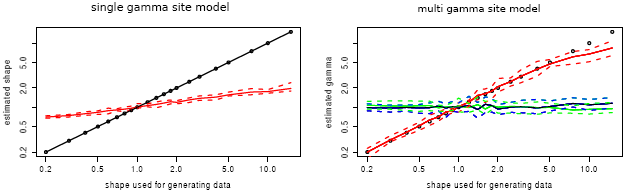
Simulated and estimated values of gamma shapes when varying a single gamma and leave the remaining at 1. Both axis are log-scale.

### 3.2 Nucleotide alignments from TreeBase

To get an impression of how the multi-gamma model performs on empirical data, we randomly selected 50 nucleotide alignments of various sizes from TreeBase [26]. Appendix A lists the details of the alignments. The analysis uses a HKY substitution model [17], the uncorrelated relaxed clock model with log normal distribution [9] and Yule tree prior [35]. The model was run both with single gamma and a category for invariant sites, and results compared with the multi-gamma and relaxed gamma model, both with a category for invariant sites. The analyses were run till convergence, judging from effective sample sizes of at least 200.

Figure 3 shows results for the 50 datasets in order of alignment size, which is the number of nucleotides contained in the alignment, that is, the number of taxa times the length of the alignment. The top shows differences in likelihood estimates and AICM [1] scores for these analyses. It appears that for smaller alignments, there is not much benefit of using the multi-gamma model. However, it does not cause much damage either. For larger alignments, there can be significant improvement of the fit. Given these data sets were randomly selected, we expect this to be true for many data sets in general.

**Figure 3.**
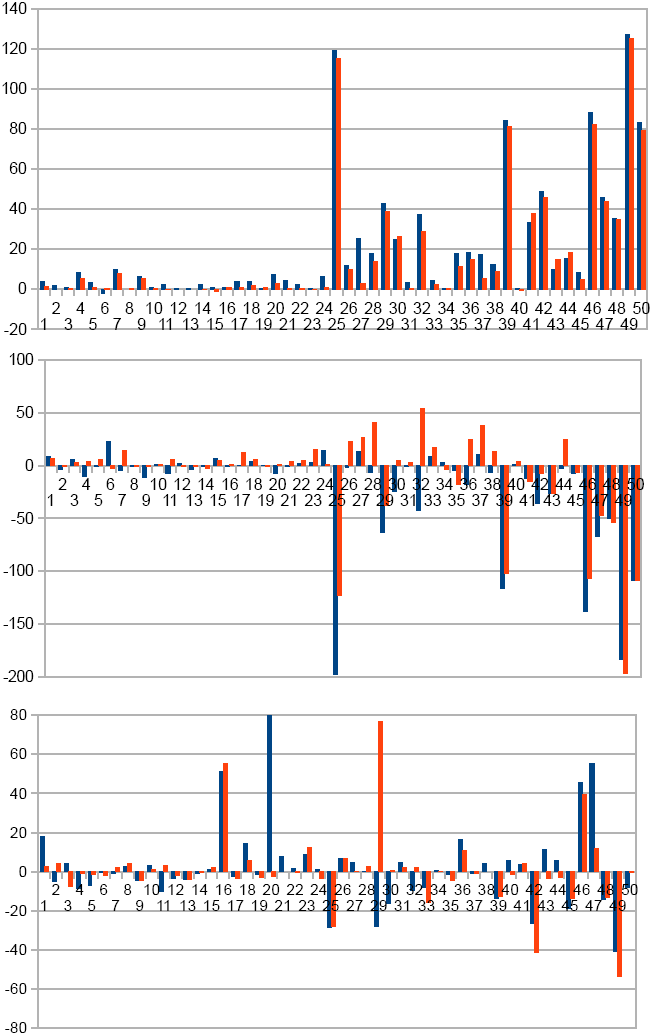
Results for 50 TreeBase alignments ordered in alignment size from small to big comparing single gamma, with multi-gamma models (in blue) and relaxed gamma models (in red). Difference in log likelihoods at the top (higher favours multi-gamma models) and difference of AICM in the middle (lower favours multi gamma models). The bottom shows percentage change in root height estimates.

The bottom of Figure 3 shows there are sometimes dramatic differences in divergence time estimates. This observation highlights the concern for the impact that model misspecification can have on molecular dating.

This conclusion is supported by similar analyses of the datasets from the partition finder repository (see Appendix D). Comparing results for single and relaxed gamma site models that assume a proportion of invariable sites, we find that for those datasets that converged (not all of them did, since some datasets were too large to converge over the period of analysis) relaxed gamma site models in general fit better for any substantial dataset, judging from AICM estimates. Further, time estimates can differ considerably between models.

### 3.3 Dating the origin of green plants and algae

We collected a set of 34 atpB and rpoC1 chloroplast genes from Genbank (see Appendix B for accession numbers), aligned their inferred protein sequences using MUSCLE [10] in MEGA [31] and then converted the protein sequences back to nucleotides. This produced data matrices for atpB with 1401 sites (749 patterns) and for rpoC1 with 1359 sites (1097 patterns). Furthermore, we constrained the tree to the topology shown in Figure 4 and we have two calibrations one on the stem leading to angiosperms following divergence from conifers and one on the stem leading to ferns subsequent to their divergence from seed plants [27] (Figure 4).

**Figure 4.**
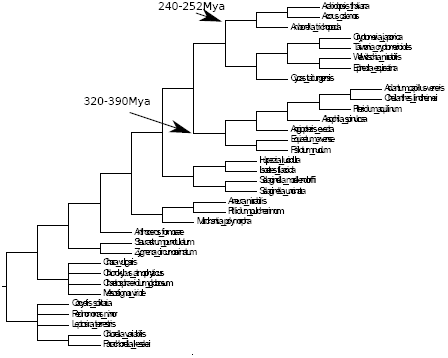
Restrictions on tree and fossil calibrations.

Data was analysed in BEAST [5] by assuming a separate partition for each codon position. A reversible-jump based substitution model [3] was used and GTR was found to be preferred in all cases, with the exception of partition 3 of the atpB dataset, for which the EVS model was preferred. A discrete four category gamma distribution model that also assumed invariable sites, was found to fit the data much better than a model that assumed no positional rate heterogeneity or a model that assumed a single rate class plus invariable sites. We used an uncorrelated relaxed clock (log normal) model [9] for branches, and estimated a coefficient of variation of around 0.7 for both genes. This result indicates that that the strict clock can be rejected. A Yule process was assumed for the tree prior. The main issue when using a single-gamma site model is that root height estimates differ considerably, with mean estimates leaving a gap of over 100M year. Also, the topology in the part of the tree that is not restricted differs between the genes. Given both genes are chloroplast genes, hence must share a common ancestry, one would expect the difference between time estimates to be much smaller, as well as showing support for the same topology. See Appendix C for details on the BEAST XML files.

Figure 5 shows the marginal likelihoods for the two genes, with single-gamma, multi-gamma and relaxed gamma models. The relaxed-gamma model was run as a single model shared by the three partitions, and a variant that has separate relaxed-gamma models for each of the three partitions. Figure 5 shows the latter model fits much better than the former as indicated by the non-overlapping densities of log likelihood, while both perform much better than the single-gamma model. The multi-gamma model applied to all three partitions (that is, three shape parameters per branch) fits similarly to the relaxed-gamma model; slightly worse for atpB and slightly better for rpoC1.

**Figure 5.**
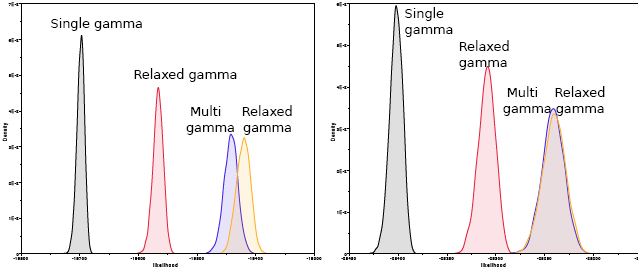
Marginal likelihood for atpB (left) and rpoC1 (right) analyses under the relaxed-gamma model.

Figure 6 shows a DensiTree [4] of the atpB and rpoC1 tree-posteriors. Switching from single to relaxed-gamma-model does not change the inferred topology very much, and the differences in topology is illustrated in the figure.

**Figure 6.**
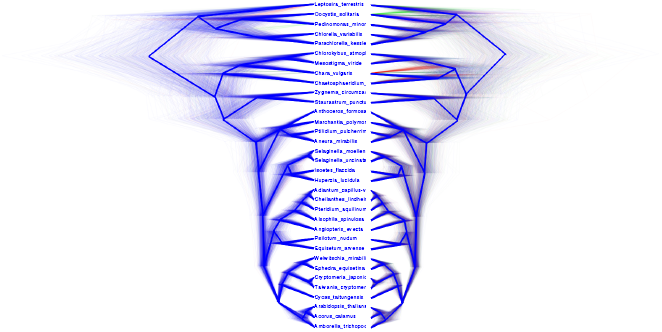
Topology under atpB (left) and rpoC1 (right)

Figure 7 shows marginal densities of root height estimates. It shows that mean root height estimates for the atpB and rpoC1 gene differ much less using the multi-gamma (79My) model than the single-gamma model (107My). Using the relaxed gamma model makes the estimates even less dramatic, reducing the difference to 28My. Note that the densities between atpB and rpoC1 overlap much more for the multi and relaxed gamma-models than the single-gamma model, indicating less discrepancy between the mdoels. However, the 95% HPDs of root heights becomes much larger under the latter models.

**Figure 7.**
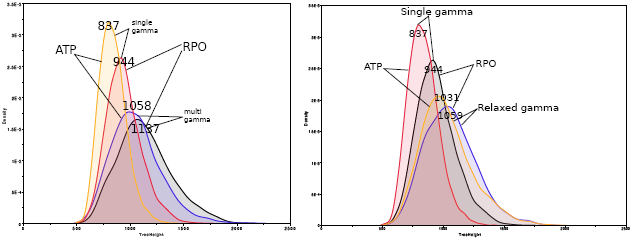
Root height estimates under the various models.

It is notable that there is a rate paradox for this data; estimates for clock rates are higher for multi-gamma and relaxed-gamma models (mean rate 6.86E-10 and 1.03E-9 substitutions per site per year for single gamma and 9.17E-10 and 1.04E-9 s/s/y for relaxed-gamma model for APT and rpoC1 respectively). The higher clock rate is consistent with a better fit, but one would expect a lower root to accommodate the mutations in the data, but the models also infer a higher root date instead of a lower one. This suggests that for this data the multi/relaxed gamma model accommodate more mutations. An alternative interpretation of our results is that the single-gamma model underestimates the amount of mutation.

The analysis was repeated with atpB and rpoC1 concatenated into a single alignment, which seems reasonable given both are from the chloroplast genome, hence must share a common history. Essentially, all results can be interpreted as the average of the results for the individual atpB and rpoC1 analyses. The fit for multi/relaxed-gamma is still much better than for single gamma, the root estimate higher, and in between that of atpB and rpoC1 individually, and topology estimate is partly compatible with the atpB topology and partly with the rpoC1 topology.

## 4 Discussion

We introduced a Bayesian framework for allowing shape parameters governing rate heterogeneity to vary across branches. The method allows us to get consistent time estimates for different chloroplast genes for green plants, while without the multi-gamma rate model we get substantially different estimates. This suggests that the heterotachy suspected for this data can be captured effectively through multi-gamma site models. Furthermore, for many datasets, the method gives a much better fit as well as different time estimates.

The method is implemented in the MGSM (multi-gamma site model) package for BEAST [5]. It is open source and freely available under LGPL license from https://github.com/BEAST2-Dev/MGSM/, and offers GUI support through BEAUti for setting up an analysis (see https://github.com/BEAST2-Dev/MGSM/wiki).

It did not escape our attention that the techniques for handling branch related parameters as in the multi-gamma and relaxed gamma models could be used for other features of substitution models over branches. Possible directions for generalising multi-gamma site models include site models with a limited number of gamma shapes, in a similar fashion as clock models such as the random local clock [7] can have a limited number of branch rates in between 1 and the total number of branches.

## Acknowledgements

This work was assisted by stimulating discussions with Chris Simon and Dave Marshall as well as comments at the New Zealand Phylogenomics 2015 meeting. This work was funded by a Rutherford fellowship (http://www.royalsociety.org.nz/programmes/funds/rutherford-discovery/) from the Royal Society of New Zealand awarded to Prof. Alexei Drummond.

## Appendix A: List of alignments from TreeBase

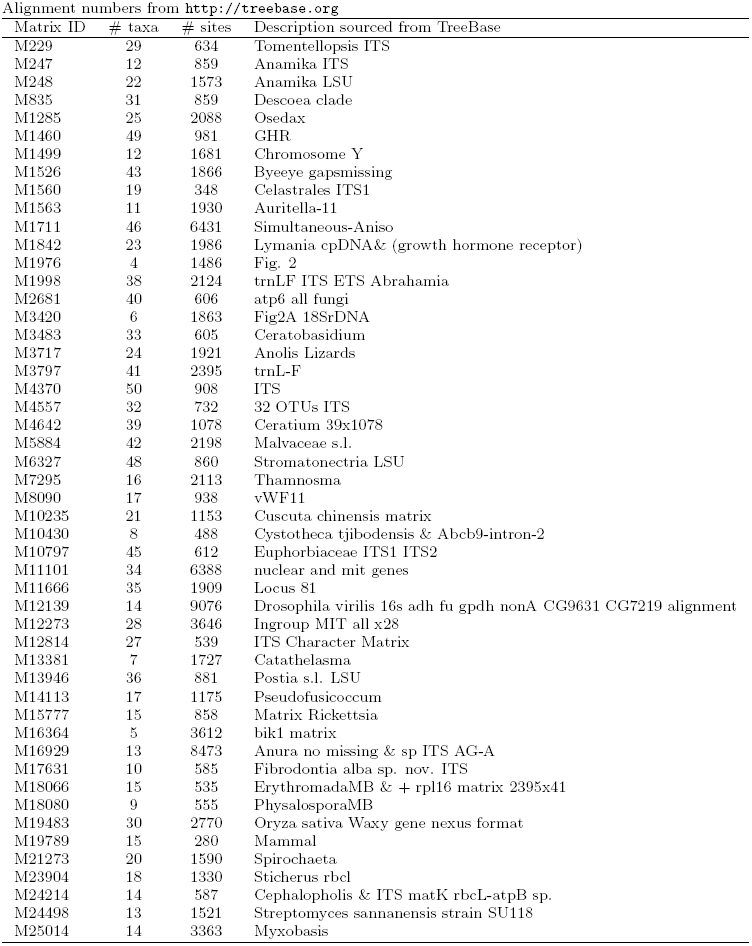

Summary table for nucleotide sequences from TreeBase. The number in first column coincides with numbers in Figure 3

|  | Dataset | 4gi |  |  | 4mgi |  |  | 4rgi |  |  |
| --- | --- | --- | --- | --- | --- | --- | --- | --- | --- | --- |
|  |  | logL | AICM | Height | logL | AICM | Height | logL | AICM | Height |
| 1 | M19789 | -2136.8475 | 4308.693 | 0.3928786266 | -2132.9903 | 4316.91 | 0.4022 | -2135.3667 | 4315.194 | 0.3495 |
| 2 | M10430 | -1204.2801 | 2425.789 | 0.1307 | -1202.565 | 2422.413 | 0.1239 | -1204.3171 | 2424.995 | 0.1361 |
| 3 | M18080 | -1950.8533 | 3919.156 | 0.1597 | -1949.9263 | 3924.941 | 0.1663 | -1950.7097 | 3921.987 | 0.1472 |
| 4 | M17631 | -3460.2928 | 6941.796 | 0.7876 | -3451.878 | 6931.547 | 0.7186 | -3454.9799 | 6945.568 | 0.7819 |
| 5 | M1976 | -3540.447 | 7088.348 | 0.1163 | -3537.0736 | 7087.692 | 0.1079 | -3539.7286 | 7093.989 | 0.1145 |
| 6 | M1560 | -3136.5259 | 6312.933 | 0.4957 | -3138.8655 | 6335.85 | 0.4928 | -3136.0007 | 6309.815 | 0.4847 |
| 7 | M18066 | -2675.0575 | 5385.319 | 0.1589 | -2665.2159 | 5380.643 | 0.1572 | -2667.068 | 5399.271 | 0.1621 |
| 8 | M24214 | -907.8032 | 1839.586 | 0.0043297 | -907.8311 | 1838.779 | 0.0044378 | -907.7367 | 1838.841 | 0.0045036 |
| 9 | M247 | -2933.511 | 5914.866 | 0.1612 | -2927.1371 | 5903.733 | 0.154 | -2928.0852 | 5913.916 | 0.1538 |
| 10 | M3420 | -3118.0111 | 6246.542 | 0.0225 | -3117.3701 | 6247.134 | 0.0232 | -3117.6888 | 6247.359 | 0.0227 |
| 11 | M15777 | -1824.6666 | 3685.85 | 0.0271 | -1822.5692 | 3678.614 | 0.0243 | -1824.8101 | 3691.549 | 0.028 |
| 12 | M13381 | -3714.5649 | 7452.41 | 0.0481 | -3714.1537 | 7454.202 | 0.0463 | -3714.6251 | 7451.283 | 0.0471 |
| 13 | M12814 | -937.2828 | 1911.745 | 0.0071562 | -936.8968 | 1908.188 | 0.0068783 | -937.2707 | 1911.323 | 0.0068832 |
| 14 | M8090 | -2738.5756 | 5508.114 | 0.0497 | -2736.4928 | 5506.812 | 0.0492 | -2738.9789 | 5504.87 | 0.0495 |
| 15 | M229 | -2187.3138 | 4429.037 | 0.0678 | -2186.4342 | 4435.671 | 0.0687 | -2188.3429 | 4434.021 | 0.0692 |
| 16 | M14113 | -2898.4699 | 5823.49 | 0.4174 | -2897.5843 | 5822.312 | 0.6306 | -2897.8555 | 5824.87 | 0.6475 |
| 17 | M3483 | -1975.2533 | 4012.721 | 0.0372 | -1971.3285 | 4012.63 | 0.0363 | -1974.4593 | 4025.041 | 0.0359 |
| 18 | M24498 | -2755.5961 | 5556.459 | 0.0176 | -2751.7179 | 5560.086 | 0.0201 | -2753.6969 | 5561.925 | 0.0186 |
| 19 | M1499 | -3125.8316 | 6273.332 | 0.0246 | -3125.6009 | 6273.723 | 0.0242 | -3125.2416 | 6272.231 | 0.0239 |
| 20 | M16364 | -13356.1059 | 26724.697 | 1.1466 | -13348.8267 | 26717.337 | 3.3584 | -13353.093 | 26725.96 | 1.1155 |
| 21 | M1563 | -5890.6771 | 11810.842 | 0.1149 | -5886.5602 | 11810.292 | 0.1235 | -5890.5648 | 11815.067 | 0.1149 |
| 22 | M23904 | -3275.5349 | 6581.852 | 0.0439 | -3273.1268 | 6583.667 | 0.0447 | -3275.3788 | 6586.233 | 0.0436 |
| 23 | M10235 | -2703.819 | 5450.485 | 0.0289 | -2703.3143 | 5453.441 | 0.0315 | -2704.2486 | 5465.889 | 0.0324 |
| 24 | M4557 | -2971.1893 | 6014.962 | 0.0907 | -2964.7763 | 6029.154 | 0.0916 | -2970.2121 | 6015.927 | 0.0875 |
| 25 | M2681 | -9846.1911 | 19803.672 | 0.9047 | -9727.154 | 19606.03 | 0.6464 | -9731.0339 | 19680.873 | 0.6493 |
| 26 | M835 | -4292.5875 | 8663.089 | 0.1482 | -4280.587 | 8660.828 | 0.1579 | -4283.0637 | 8685.92 | 0.1585 |
| 27 | M10797 | -8018.9627 | 16144.199 | 0.475 | -7993.7406 | 16157.387 | 0.4966 | -8016.2651 | 16170.728 | 0.4762 |
| 28 | M13946 | -4078.8918 | 8226.82 | 0.0736 | -4061.1762 | 8220.067 | 0.0735 | -4065.0459 | 8267.309 | 0.0754 |
| 29 | M21273 | -9533.7946 | 19111.588 | 0.2792 | -9490.9268 | 19048.554 | 0.2012 | -9495.1618 | 19073.77 | 0.4941 |
| 30 | M248 | -4210.2705 | 8482.219 | 0.075 | -4185.5152 | 8457.427 | 0.0626 | -4183.9474 | 8487.001 | 0.0753 |
| 31 | M7295 | -4694.4796 | 9413.209 | 0.0526 | -4691.2445 | 9412.467 | 0.0552 | -4693.9381 | 9416.621 | 0.0536 |
| 32 | M6327 | -4813.2593 | 9774.09 | 0.0723 | -4776.1245 | 9731.539 | 0.0653 | -4784.6078 | 9827.737 | 0.0738 |
| 33 | M4642 | -2707.7318 | 5481.546 | 0.0227 | -2703.4576 | 5489.916 | 0.0209 | -2705.4797 | 5498.721 | 0.0191 |
| 34 | M1842 | -3110.2733 | 6270.127 | 0.0070593 | -3109.9808 | 6273.357 | 0.0070998 | -3110.0356 | 6266.374 | 0.0070534 |
| 35 | M3717 | -21781.2882 | 43634.435 | 0.5391 | -21763.6884 | 43629.967 | 0.5323 | -21770.0847 | 43616.439 | 0.5159 |
| 36 | M4370 | -4529.6939 | 9159.546 | 0.1538 | -4511.5996 | 9141.707 | 0.1788 | -4515.0267 | 9184.345 | 0.17 |
| 37 | M1460 | -6946.0161 | 14019.666 | 0.0792 | -6928.8923 | 14030.192 | 0.0786 | -6940.5631 | 14057.9 | 0.0784 |
| 38 | M25014 | -7314.2152 | 14662.746 | 0.0453 | -7302.0321 | 14656.572 | 0.0472 | -7305.4891 | 14675.943 | 0.0453 |
| 39 | M1285 | -13355.6276 | 26766.856 | 0.3152 | -13271.5138 | 26650.638 | 0.2716 | -13274.5577 | 26664.298 | 0.2751 |
| 40 | M11666 | -2911.0707 | 5853.877 | 0.0017217 | -2910.846 | 5854.405 | 0.0018171 | -2911.5679 | 5858.014 | 0.0016958 |
| 41 | M1526 | -5664.4201 | 11453.866 | 0.024 | -5631.2512 | 11441.508 | 0.0249 | -5626.6066 | 11439.015 | 0.025 |
| 42 | M1998 | -11564.6916 | 23200.15 | 0.1683 | -11515.8625 | 23163.982 | 0.1239 | -11518.8415 | 23192.211 | 0.0984 |
| 43 | M19483 | -4708.4409 | 9486.533 | 0.0113 | -4698.736 | 9460.552 | 0.0126 | -4693.856 | 9460.117 | 0.0109 |
| 44 | M5884 | -6898.4465 | 13882.141 | 0.0277 | -6882.9889 | 13879.288 | 0.0293 | -6880.2777 | 13906.713 | 0.0268 |
| 45 | M3797 | -5973.4118 | 12052.914 | 0.0208 | -5965.0801 | 12045.769 | 0.0169 | -5968.6401 | 12046.771 | 0.0179 |
| 46 | M12273 | -22793.8586 | 45669.023 | 0.1936 | -22705.5187 | 45530.638 | 0.2818 | -22711.8045 | 45561.898 | 0.2701 |
| 47 | M16929 | -29828.8968 | 59687.073 | 0.0706 | -29782.9779 | 59619.427 | 0.1097 | -29785.2618 | 59640.032 | 0.0789 |
| 48 | M12139 | -29070.5222 | 58173.63 | 0.1259 | -29035.189 | 58123.616 | 0.1082 | -29035.8247 | 58119.319 | 0.109 |
| 49 | M11101 | -47426.5181 | 94952.268 | 0.3398 | -47299.4892 | 94768.779 | 0.2003 | -47301.207 | 94755.571 | 0.1572 |
| 50 | M1711 | -38627.0072 | 77368.97 | 0.1044 | -38543.7533 | 77260.275 | 0.0957 | -38547.8281 | 77260.311 | 0.1038 |

## Appendix B: Genbank sequences

| Name | Genbank accession number |
| --- | --- |
| *Arabidopsis thaliana* | NC_000932 |
| *Acorus calamus* | NC_007407 |
| *Amborella trichopoda* | NC_005086 |
| *Cryptomeria japonica* | NC_010548 |
| *Taiwania cryptomerioides* | NC_016065 |
| *Cycas taitungensis* | NC_009618 |
| *Welwitschia mirabilis* | NC_010654 |
| *Ephedra equisetina* | NC_011954 |
| *Anthoceros formosae* | NC_004543 |
| *Syntrichia ruralis* | NC_012052 |
| *Chara vulgaris* | NC_008097 |
| *Chlorokybus atmophyticus* | NC_008822 |
| *Chaetosphaeridium globosum* | NC_004115 |
| *Huperzia lucidula* | NC_006861 |
| *Isoetes flaccida* | NC_014675 |
| *Selaginella moellendorffii* | NC_013086 |
| *Selaginella uncinata* | AB197035 |
| *Aneura mirabilis* | NC_010359 |
| *Marchantia polymorpha* | NC_001319 |
| *Ptilidium pulcherrimum* | NC_015402 |
| *Mesostigma viride* | NC_002186 |
| *Adiantum capillus-veneris* | NC_004766 |
| *Alsophila spinulosa* | NC_012818 |
| *Angiopteris evecta* | NC_008829 |
| *Cheilanthes lindheimeri* | NC_014592 |
| *Equisetum arvense* | NC_014699 |
| *Psilotum nudum* | NC_003386 |
| *Pteridium aquilinum* | NC_014348 |
| *Staurostrum punctulatum* | NC_008116 |
| *Zygnema circumcarinatum* | NC_008117 |
| *Chlorella variabilis* | NC_015359 |
| *Oocystis solitaria* | FJ968739 |
| *Parachlorella kessleri* | NC_012978 |
| *Pedinomonas minor* | NC_016733 |
| *Leptosira terrestris* | NC_009681 |

## Appendix C: XML for green plants and algae analysis

All BEAST XML files can be downloaded from here: https://www.cs.auckland.ac.nz/~remco/MGSMxml.tgz.

All analyses uses partitions for each 3 codon positions, a reversible-jump based substitution model, uncorrelated log normal clock model and Yule tree prior.

atpB = atpB chloroplast gene from green plants and algae

rpoc = rpoC1 chloroplast gene from green plants and algae

atprpo = atpB and rpoC1 genes concatenated

|  |  |
| --- | --- |
| atpB.RB4g.ucln.yule.xml | single gamma site categories, one gamma per partition |
| atpB.RB4spmg.ucln.yule.xml | multi gamma site model, one gamma per branch (shared among partitions) |
| atpB.RB8spmg.ucln.yule.xml | " |
| atpB.RB4mg.ucln.yule.xml | multi gamma site model, one gamma per partition per branch |
| atpB.RB8mg.ucln.yule.xml | " |
| atpB.RB4gi.ucln.yule.xml | single gamma site categories, one gamma per partition + invariant sites |
| atpB.RB4mgi.ucln.yule.xml | multi gamma site model, one gamma per branch per partition + invariant sites |
| atpB.RB4rmgi.ucln.yule.xml | relaxed gamma site model, one gamma per branch + invariant sites |
| atpB.RB43rmgi.ucln.yule.xml | relaxed gamma site model, one gamma per branch per partition + invariant sites |
| rpoc.RB4g.ucln.yule.xml | single gamma site categories, one gamma per partition |
| rpoc.RB4spmg.ucln.yule.xml | multi gamma site model, one gamma per branch (shared among partitions) |
| rpoc.RB8spmg.ucln.yule.xml | " |
| rpoc.RB4mg.ucln.yule.xml | multi gamma site model, one gamma per partition per branch |
| rpoc.RB8mg.ucln.yule.xml | " |
| rpoc.RB4gi.ucln.yule.xml | single gamma site categories, one gamma per partition + invariant sites |
| rpoc.RB4mgi.ucln.yule.xml | multi gamma site model, one gamma per branch per partition + invariant sites |
| rpoc.RB4rmgi.ucln.yule.xml | relaxed gamma site model, one gamma per branch + invariant sites |
| rpoc.RB43rmgi.ucln.yule.xml | relaxed gamma site model, one gamma per branch per partition + invariant sites |
| atprpo.RB4gi.ucln.yule.xml | single gamma site categories, one gamma per partition + invariant sites |
| atprpo.RB4mgi.ucln.yule.xml | multi gamma site model, one gamma per branch per partition + invariant sites |

## Appendix D: Results for partition finder repository data

Appendix D: Results for partition finder repository data Comparing single and multi-gamma site models.
|  | Delta logL | Delta AICM | Height | min(ESS) |
| --- | --- | --- | --- | --- |
| Bergsten2013 | 60.3459 | -104.168 | -18.4345281639 | 233.9029 |
| Caterino2001 | 54.834 | -26.444 | 10.6810387065 | 103.5102 |
| Delsuc2003 | 25.1585 | -33.03 | -2.9676258993 | 249.2825 |
| Dsouli2011 | 58.7006 | -60.973 | -8.1706435286 | 120.4567 |
| Endicott2008 | 42.0513 | -14.217 | 1.7730404264 | 107.0668 |
| Fishbein2001 | 79.6407 | -81.564 | 5.3448275862 | 102.6875 |
| Kang2013b | 24.7453 | -23.423 | -8.867427568 | 133.3271 |
| Sauquet2011 | 60.853 | -83.947 | -6.3829787234 | 97.4762 |

Other datasets did not get to convergence.

